# A Nanoluciferase biosensor to investigate endogenous chemokine secretion and receptor binding

**DOI:** 10.1101/2020.08.19.257469

**Authors:** Carl W. White, Kevin D. G. Pfleger, Stephen J. Hill

## Abstract

Secreted chemokines are critical mediators of cellular communication that elicit intracellular signalling by binding membrane-bound receptors. Here we demonstrate the development and use of a sensitive real-time approach to quantify secretion and receptor binding of native chemokines in live cells to better understand their molecular interactions and function. CRISPR/Cas9 genome-editing was used to tag the chemokine CXCL12 with the Nanoluciferase fragment HiBiT. CXCL12 secretion was subsequently monitored and quantified by luminescence output. Binding of tagged CXCL12 to either chemokine receptors or membrane glycosaminoglycans could be monitored due to the steric constraints of Nanoluciferase complementation. Furthermore, binding of native CXCL12-HiBiT to AlexaFluor488-tagged CXCR4 chemokine receptors could also be distinguished from glycosaminoglycan binding and pharmacologically analysed using BRET. These live cell approaches combine the sensitivity of Nanoluciferase with CRISPR/Cas9 genome-editing to detect, quantify and monitor binding of low levels of native secreted proteins in real time.

## Introduction

Secreted peptides and proteins are critical for cellular communication, with changes in expression, secretion and subsequent binding to their cellular targets mediating numerous (patho)-physiological cellular responses. Chemokines are a family of small cytokines secreted by cells that bind and activate G protein-coupled receptors, resulting in: immune cell migration; cell differentiation and development; and cancer metastasis^1^. The function of chemokines is controlled at the transcriptional as well as the post-translational level. For example, numerous factors are known *in vivo* to induce the expression and production of chemokines to recruit immune cells to inflamed tissue, while secreted chemokines bind membrane-bound proteoglycans to form chemotactic gradients which guide immune cell migration^2,3^. Investigating and quantifying chemokine ligand secretion from cells, as well as interactions with glycosaminoglycans (GAGs) found on proteoglycans and their receptors, is therefore important to properly understand chemokine regulation and function.

CXCL12, also known as stromal derived factor, is a prototypical chemokine that binds CXCR4 to mediate immune cell migration and cellular differentiation^4^, and is a known biomarker for a number of cancers^5^. Like many secreted proteins, methods to quantify chemokine expression rely on monitoring mRNA transcript levels, or mass spectrometry and immunoassay assays such ELISA and Western blotting to determine protein levels. However, these methods have limited sensitivity for detecting poorly expressed proteins and, in the case of immunoassays, rely on the availability of sensitive and selective antibodies^6^. Such approaches are also relatively low throughput and lack the temporal fidelity required to investigate chemokine secretion in a real-time manner. Additionally, these methods only inform on the expression of the chemokine without imparting knowledge of downstream signalling. Detailed pharmacological analysis of chemokine binding to their cellular targets, i.e. chemokine receptors, proteoglycans or GAGs, is therefore commonly monitored separately using approaches such as radio- or fluorescent ligand binding^7^ and surface plasmon resonance assays^8^. Such assays performed in live cell formats may struggle to differentiate between the different binding sites, whereas those configured to look at specific interactions, e.g. surface plasmon resonance, are performed with purified receptor and therefore occur in more artificial environments^8^. Furthermore, exogenous chemokine is used rather than chemokines secreted from cells and expressed under endogenous promotion.

Fusion of a luciferase to a protein of interest has allowed a wide range of biological effects to be investigated by bioluminescent technologies such as luciferase complementation and bioluminescence resonance energy transfer (BRET)^9^. Luciferase reporters have high signal-to-noise ratios, therefore providing highly sensitive detection of low abundance proteins as well as excellent quantitation over an extensive linear concentration range^10,11^. Previously, bioluminescence approaches using full length or split Gaussia luciferase have been used to investigate CXCL12 binding to CXCR4 and to the atypical chemokine receptor ACKR3 both *in vitro* and *in vivo*^12–14^. However, these studies used either purified luciferase-tagged CXCL12 or cells secreting exogenous luciferase-tagged CXCL12 to monitor binding by luciferase complementation or changes in luminescence, rather than endogenously expressed CXCL12. More recently, we have demonstrated that ligand binding to CXCR4 and ACKR3 tagged with the Nanoluciferase (NLuc) can be monitored in live cells using CXCL12 labelled with AF647 and NanoBRET^15^. It has been shown that a split-version of NLuc can be used to investigate peptide ligand binding to relaxin peptide family receptor 3 and 4 by NLuc complementation^16^. However, in these studies the peptide used to monitor these interactions was exogenously derived.

Taking advantage of the brightness of NLuc^11^, we and others have used CRISPR/Cas9-mediated genome engineering to tag and then study genes and proteins expressed under endogenous promotion using either full length or split NLuc. This has allowed changes in gene expression or protein levels to be measured and quantified^17–19^, as well as ligand binding and internalisation^15,20^, protein-protein interactions^15,21^, post-translational modifications^19^, and protein degradation^22^ to be monitored in real time live cell or lysed cell assays by NanoBRET, or changes in luminescence for NLuc complementation. While these studies using CRISPR/Cas9 genome-editing and luciferase tags have principally investigated membrane bound and intracellular proteins, Vascular Endothelial Growth Factor (VEGFA), a secreted growth factor, was previously tagged with the small NLuc fragment (HiBiT)^19^ and expression measured in a lytic cell assay at a single time point, indicating the potential for use of this approach to monitor secreted chemokines. Biosensors capable of investigating both ligand secretion and ligand binding of endogenously expressed proteins will be highly useful to understand cellular signalling. Using CRISPR/Cas9-mediated genome engineering, here we report a live cell assay that can be used to monitor and quantify protein secretion by luciferase complementation as well as chemokine ligand binding by BRET in real-time and in live cells.

## Results and Discussion

### Genome editing of CXCL12

CXCL12 is an essential chemokine secreted endogenously by many cells including the immortalised HEK293 cell line that is commonly used to investigate receptor pharmacology. However, real-time live cell assays to quantify endogenous CXCL12 levels are lacking. Here, to monitor expression and secretion of endogenous CXCL12 from HEK293 cells we first used CRISPR/Cas9 genome-editing to append the small 11 amino acid fragment of NLuc, HiBiT, to the C-terminus of CXCL12. The use of a small Nluc fragment rather than full length NLuc simplifies and improves the efficiency of the genome-editing process in cells. Indeed, we found homozygous HiBiT insertion into the native *CXCL12* locus (Supplementary Figure 1), a low probability event in non-diploid HEK293 cells. The minimal size also reduces the potential of the tag to perturb the function of CXCL12, with the placement of the HiBiT tag on the C-terminus of CXCL12 based on the knowledge that the N-terminus is critical for CXCR4 activation^25^ and previous fusion of Gaussia luciferase to CXCL12 in over-expression models that did not impede function^13^. This strategy using HiBiT rather than full length NLuc also limits the observable signal to secreted (extracellular) CXCL12 due to the cell impermeant nature of the 18 kDa NLuc fragment (LgBiT) used for luciferase complementation.

### Quantification of ligand secretion

In live HEK293 cells expressing genome-edited CXCL12-HiBiT incubated with LgBiT (30 nM), we observed a gradual increase in luminescence (Figure 1a) over the course of one hour and increasing either cell number or incubation time further augmented the increase in luminescence (Figure 1a and 1b) indicating continuous CXCL12-HiBiT secretion. These results further demonstrate the applicability of genome-editing approaches to monitor protein levels, now including secreted proteins, in real-time which overcomes the low throughput non-live cell limitations of common techniques used to quantify these, such as ELISA and western blotting. However, our initial data (Figure 1) report relative changes in expression rather than provide absolute quantification of secreted protein. Such measures can be made using immunoassays and are important for pharmacological evaluation of ligand function. To address this, we sought to quantify expression through correlation with luminescent output.

**Figure 1:**
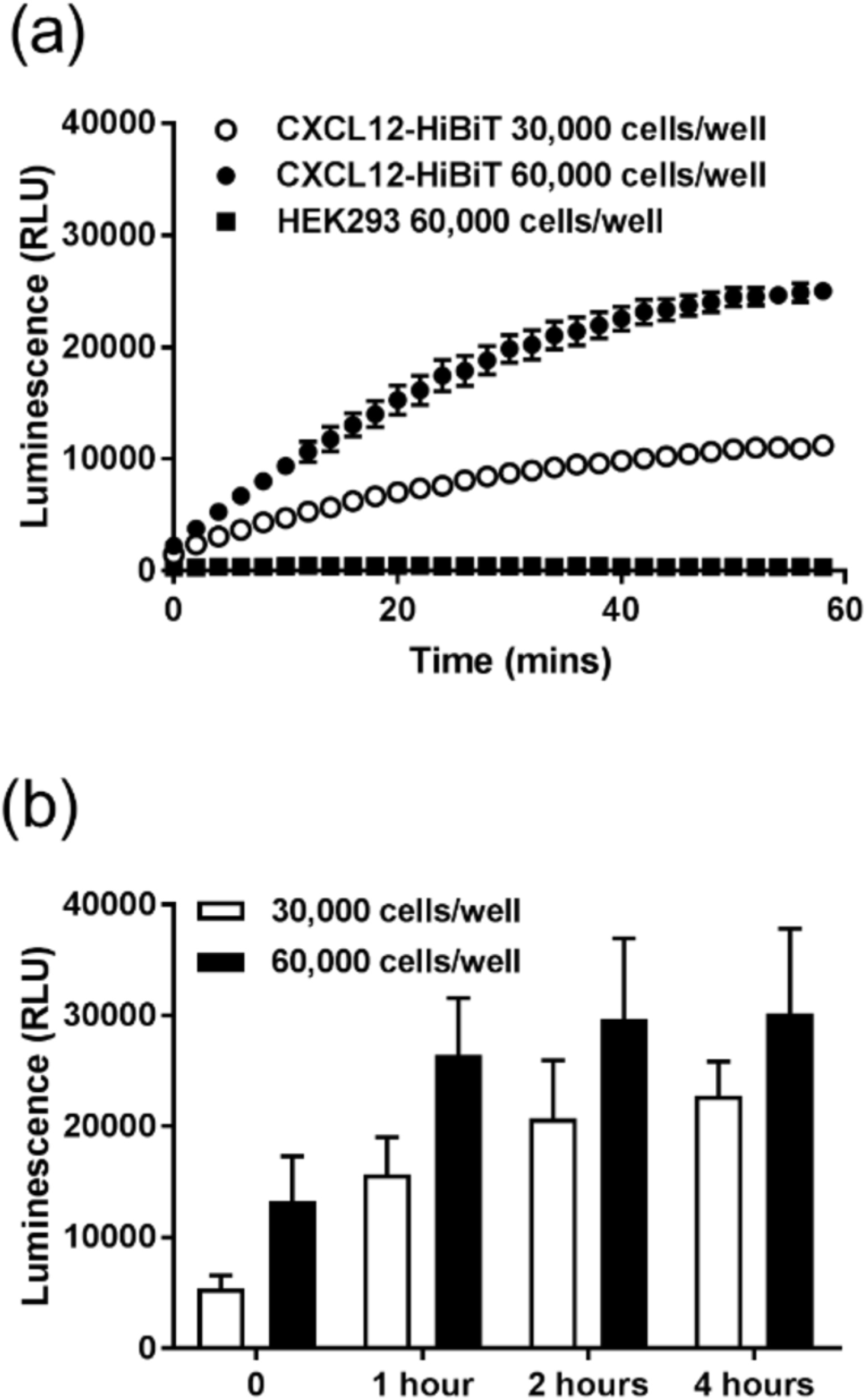
Investigation of CXCL12-HiBiT secretion from genome-edited HEK293 cells. (a) Kinetic analysis of luminescence generated post-wash from HEK293 cells expressing genome-edited CXCL12-HiBiT plated at 30,000 (white circles) or 60,000 (black circles) cells/well or wildtype HEK293 cells (squares) immediately following LgBiT (30 nM) addition. (b) Effect of cell number and post-wash incubation time on luminescence generated by addition of LgBiT (30 nM) to HEK293 cells expressing genome-edited CXCL12-HiBiT. Points and bars are (b) mean ± s.e.m. of three experiments performed in triplicate or are (a) representative of three experiments.

Previous work has shown that complementation of the two NLuc fragments (HiBiT with LgBiT) is linear, with luminescence extending over 8 orders of magnitude^19^. Here we took advantage of this relationship to quantify secreted CXCL12-HiBiT expression from our genome-edited live HEK293 cells (Figure 2). At two hours post-wash, when CXCL12 secretion had largely plateaued, we observed CXCL12-HiBiT expression to be 0.41 ± 0.10 pM/well (n=6) and 0.79 ± 0.14 pM/well (n=6) of a 96 well plate seeded with 30,000 and 60,000 cells/well respectively (Figure 2). Although the current data demonstrate absolute quantification, this is of luciferase-tagged CXCL12 under endogenous promotion rather than endogenous CXCL12 and CRISPR/Cas9 tagging of native proteins may alter expression^15,26^. While no such effects were seen previously with insertion of small NLuc tags^15^, changes in expression relative to wildtype cells expressing untagged CXCL12 due to tagging or genetic rewiring following prolonged passage may need to be considered. Commercial immunoassays for CXCL12 detection regularly have a sensitivity of >1 pg/ml (∼0.1 nM) and normal working ranges of 10-1000 pg/ml. In contrast here, the sensitivity of the luminescent approach was approximately 1000-fold greater with measurable signal in the low fM range. Moreover, we observed luminescence output that was linear over seven orders of magnitude which exceeds the typical dynamic range (2-3 orders of magnitude) seen with standard immunoassays.

**Figure 2:**
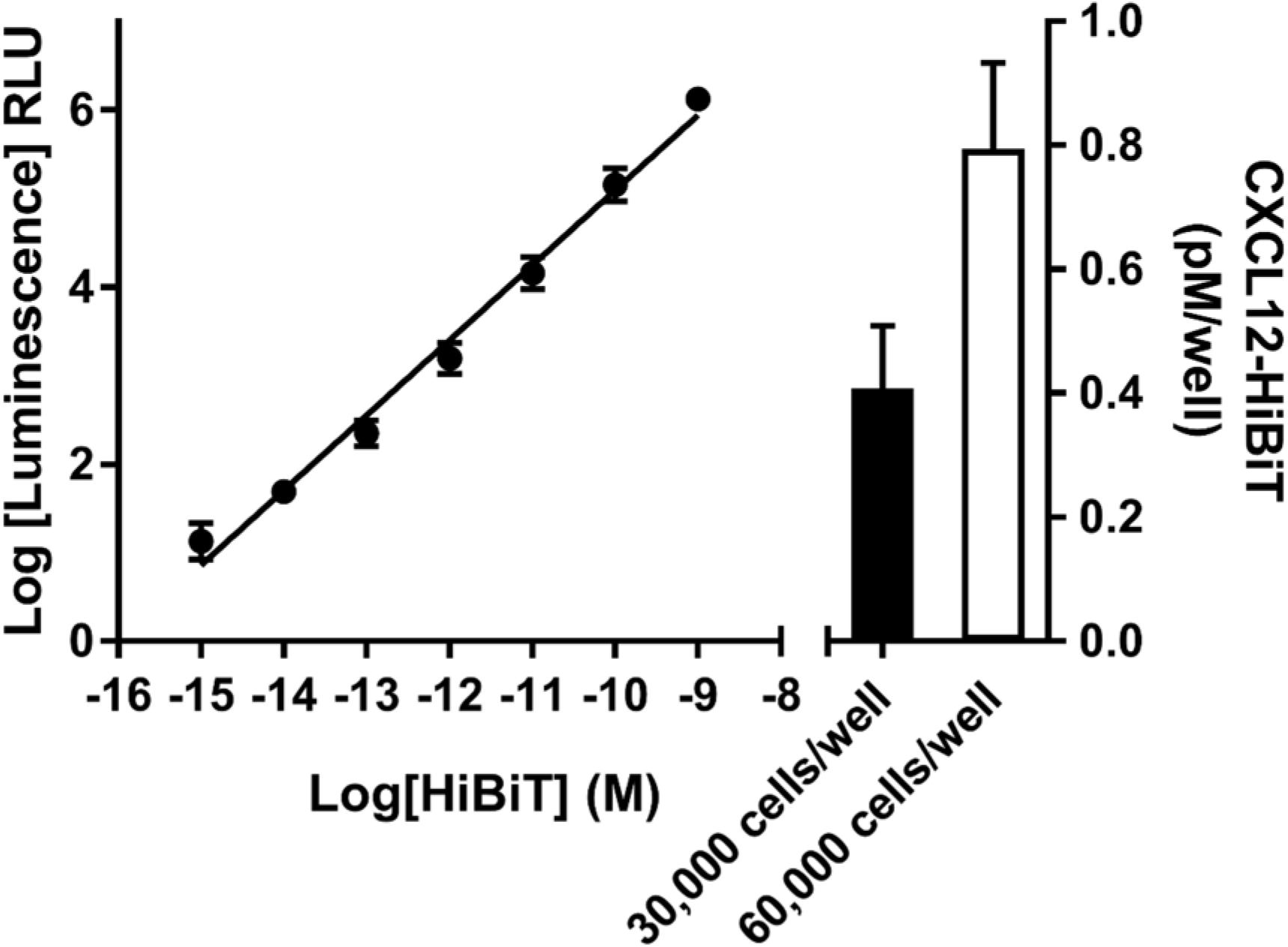
Quantification of CXCL12-HiBiT expression in genome-edited HEK293 cells. Luminescence generated from purified LgBiT (100 nM) incubated with increasing concentrations of purified HiBiT-Halotag (10 fmol - 100 nM) was used to construct a linear standard curve to quantify CXCL12-HiBiT expression by linear regression in wells seeded with either 30,000 (black bar) or 60,000 (white bar) genome-edited HEK293 cells. Points and bars are mean ± s.e.m. of six experiments performed in triplicate.

### NanoBRET ligand binding

Exogenous CXCL12 tagged with full length Gaussia-luciferase have been used to investigate ligand binding at CXCR4 and ACKR3, with fusion of the luciferase having limited effects on CXCL12 function^13^. To ensure that CXCL12 tagged with HiBiT retained functionality, we established a new NanoBRET ligand binding configuration that should be widely applicable to chemokines and other secreted proteins such as growth factors. Here, HEK293 cells expressing genome-edited CXCL12-HiBiT were co-cultured with wildtype HEK293 cells or HEK293 cells transiently transfected with SNAP/CXCR4. Binding of CXCL12-HiBiT to CXCR4 brings the donor luciferase (once it has been complimented with cell impermeant purified LgBiT) on the ligand and a fluorescent reporter on the receptor into close proximity, thereby increasing the BRET ratio that can be measured and inferred as ligand binding. Compared to coincubation with wildtype HEK293 cells, we observed an increase in BRET between CXCL12-HiBiT (complemented with purified LgBiT) and SNAP/CXCR4 labelled with cell impermeant AlexaFluor488 that was displaced in a concentration-dependent manner by the CXCR4 antagonist AMD3100 at the anticipated affinity (Figure 3, pIC_50_ = 6.97 ± 0.10, n=5). This demonstrates that the genome-edited CXCL12-HiBiT is both secreted and capable of binding to CXCR4 exogenously expressed in neighbouring cells.

**Figure 3:**
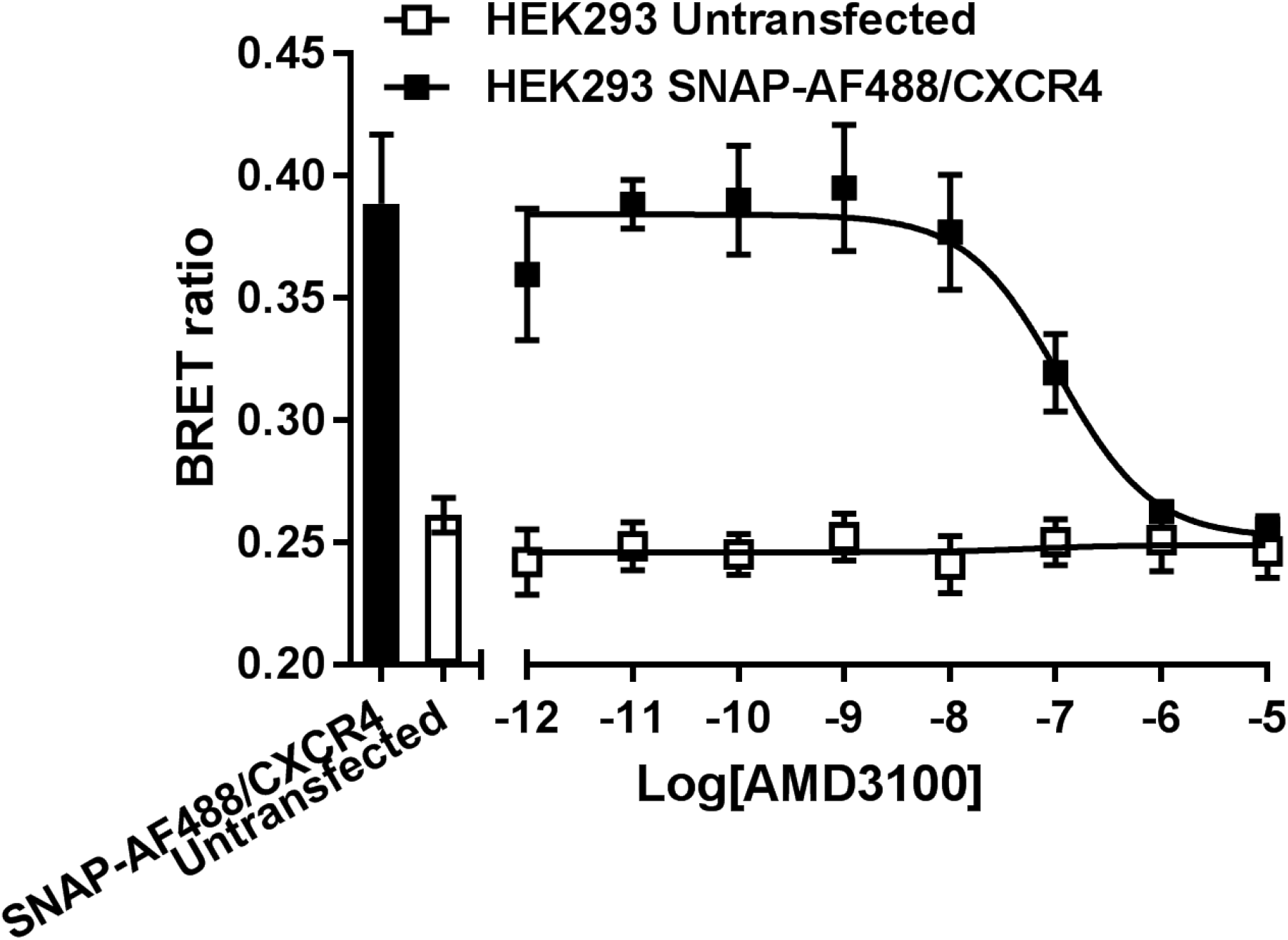
Displacement of genome-edited CXCL12-HiBiT binding to SNAP/CXCR4 observed by NanoBRET. HEK293 cells expressing genome-edited CXCL12-HiBiT, co-cultured with wildtype HEK293 cells (white squares) or HEK293 cells transiently transfected with SNAP/CXCR4 (black squares), were incubated in the absence or presence of increasing concentrations of AMD3100 (100 pM – 10 µM). CXCL12-HiBiT was complemented with purified LgBiT (30 nM) and SNAP/CXCR4 labelled with cell impermeant AlexaFluor488. Bars represent basal BRET in the absence of AMD3100. Bars and points represent mean ± s.e.m. of five individual experiments performed in duplicate.

In addition to investigating binding using CXCL12 tagged with full-length luciferase, purified or secreted CXCL12 tagged with split-Gaussia luciferase has been used to probe CXCR4 or ACKR3 tagged with the corresponding Gaussia luciferase fragment *in vitro* and *in vivo*^12^. More recent studies have used split-NLuc complementation to investigate ligand binding at the relaxin-3 receptor^16^ or binding of a Nanobody (VUN400-HiBiT) to CXCR4 tagged with split-NLuc fragments^27^. However, here we used NanoBRET rather than using luciferase complementation to investigate ligand binding. NanoBRET confers the advantage of reducing any confounding effects due to the affinity of the luciferase complementation, particularly if investigating low affinity ligand-receptor interactions. Moreover, the high distance dependence of energy transfer in NanoBRET assays distinguishes binding of CXCL12 at receptors from non-receptor binding such as at glycosaminoglycans (GAGs). Indeed, as discussed below, this may be a useful consideration since interactions with GAGs both regulate and add complexity to chemokine signalling^2^. Finally, traditional NanoBRET binding assays where the receptor is tagged with NLuc requires suitable fluorescent ligands that are not always readily available. Our approach achieves the benefits of NanoBRET ligand binding without the need to generate complex fluorescent proteins/probes that may require recombinant production.

### Detection of chemokine-GAG interactions

We have previously reported that knockout of CXCL12 from HEK293 cells decreases constitutive CXCR4 internalisation suggesting that sufficient endogenous CXCL12 is secreted to activate and internalise CXCR4^15^. However the apparent concentration of CXCL12-HiBiT that we observed is approximately 300-1000-fold lower than the reported EC_50_ for CXCL12-mediated CXCR4 G protein signalling^28^. It is known that ligand concentration within a plate based cellular assay is unlikely uniform^29^ and that secreted chemokines can be concentrated/localised at the cell surface by binding to GAGs, which increases the effective ligand concentration near the receptor^2^. We have shown previously that when NLuc fragments are fused to proteins steric constraints can modulate the affinity of complementation^15^, while binding of chemokines to GAGs facilitates oligomerisation as well as clustering^2^. We therefore hypothesised that CXCL12 binding to GAGs would impart such constraints on NLuc complementation and provide a mechanism to determine if GAG-mediated membrane accumulation was indeed occurring.

To test this, we first determined the affinity of NLuc complementation in our HEK293 cells expressing genome-edited CXCL12-HiBiT. We found that in live cells the affinity of NLuc complementation (CXCL12-HiBiT with LgBiT) was reduced (K_d_ =11.6 ± 1.48 nM, mean ± s.e.m., Figure 4a) compared to the reported affinity (∼700 pM) for purified NLuc fragments^10^ indicating that fusion of HiBiT to CXCL12 modulated complementation. Next we used a small molecule glycosaminoglycan inhibitor^30^ surfen (10 µM) to pharmacologically disrupt CXCL12-GAG binding and observed a small increase in the affinity of CXCL12-HiBiT-LgBiT complementation compared to the vehicle control (Figure 4a; K_d_ =5.93 ± 0.53 nM, mean ± s.e.m., n=6, p<0.05), as well as an increase in the luminescence output (Figure 4a and 4b). In contrast, incubation with exogenous heparan sulfate (30 µg/ml), a major glycosaminoglycan, decreased luminescence and attenuated the surfen mediated responses (Figure 4b, p<0.01, n=5). These data indicate that the differences in complementation affinity between HiBiT and LgBiT were due to conformational differences of CXCL12-HiBiT when bound to GAGs and/or in oligomeric forms compared to when found in a free state. Finally, in a kinetic analysis (Figure 4b) we also observed that incubation of cells with AMD3100 (1 µM) increased luminescence output, indicating that CXCL12-HiBiT binds to endogenous CXCR4 expressed in HEK293 cells. Combined, these results indicated that CXCL12 – GAG interactions are occurring in HEK293 cells and suggest a possible mechanism by which a cell may achieve sufficient accumulation of endogenous CXCL12 at the membrane to activate CXCR4. It is noteworthy that while HEK293 cells are a common model cell line used to study receptor function, the concurrent expression, membrane accumulation and receptor activation of endogenously-secreted ligands from these cells are under appreciated and may confound the interpretation of results.

**Figure 4:**
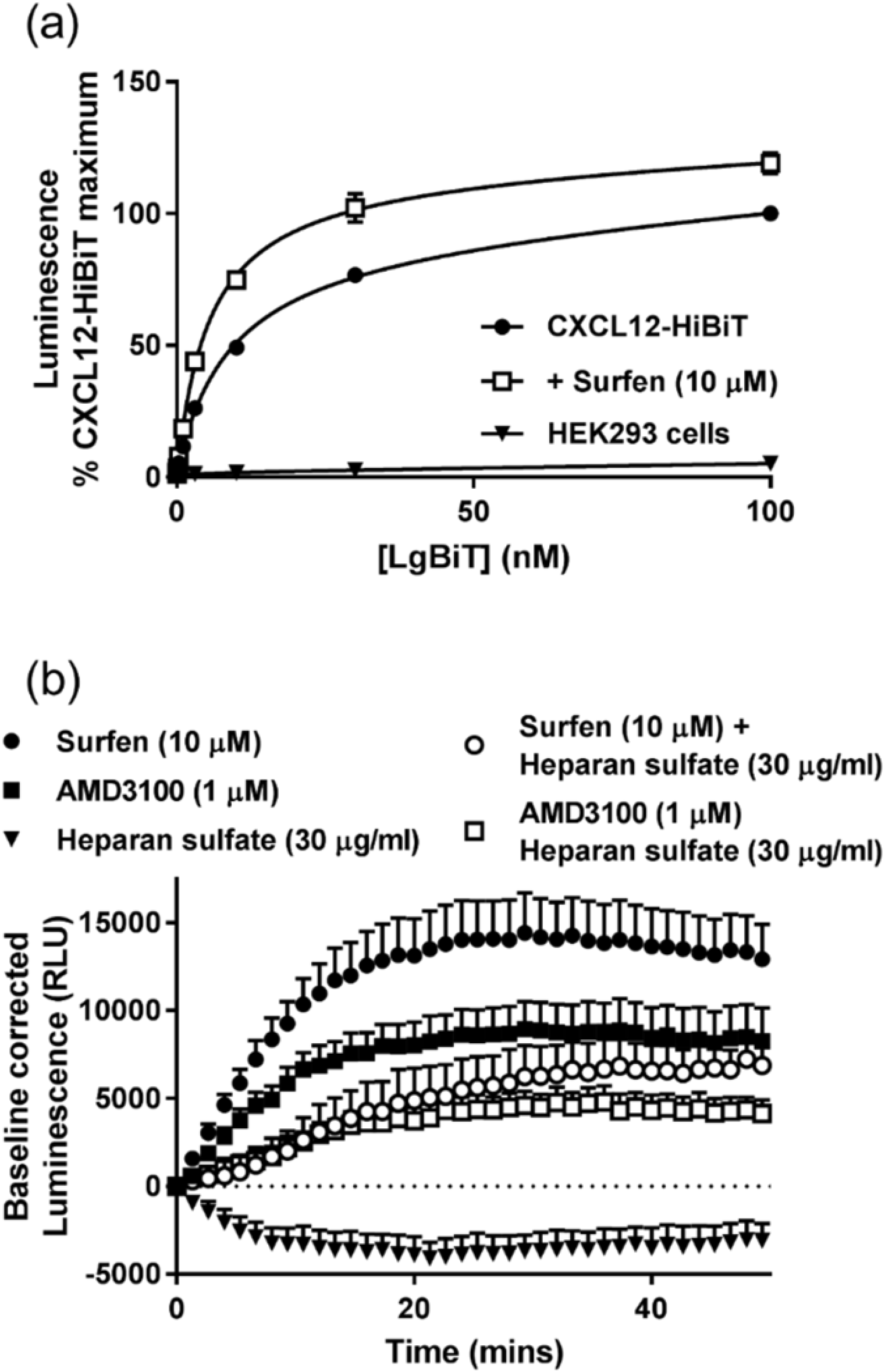
Monitoring CXCL12-glycosaminoglycan interactions by changes in luminescent output. (a) HEK293 cells expressing genome-edited CXCL12-HiBiT or wildtype HEK293 cells (downward triangles) were incubated with increasing concentrations of purified LgBiT in the absence (black circles) or presence of Surfen (10 µM, white squares). (b) Kinetic analysis of the effect of AMD3100 (1 µM, black square), surfen (10 µM, black circle), heparin sulfate (30 mg/mL, downward triangle), AMD3100 (1 µM) plus heparin sulfate (30 mg/mL, white square) or surfen (10 µM) plus heparin sulfate (30 mg/mL, white circle) on the baseline-corrected luminescence generated by HEK293 cells expressing genome-edited CXCL12-HiBiT in the presence of 30 nM LgBiT. (a) Points represent % of the maximum luminescence ± s.e.m. generated by complementation of genome-edited CXCL12-HiBiT and 100 nM LgBiT in the absence of surfen from six individual experiments performed in duplicate. (b) Points represent mean ± s.e.m. of five individual experiments performed in duplicate.

### Summary and future directions

In summary, the genome-editing approach described here allows the secretion and binding of endogenous chemokines to GAGs or receptors to be readily inferred in live cell assays through changes in luminescence and/or NanoBRET ligand binding assays. These approaches monitor continuous peptide secretion in real time and do not require the development of a selective and/or specific antibody. Importantly, binding specificity can be imparted using the NanoBRET modality that allows binding of secreted ligand to receptors to be monitored while effectively eliminating observable ‘off-target’ binding to GAGs. Like many peptides, chemokine binding to GAGs *in vivo* is vital for modulating function, however thus far these interactions have primarily been explored in artificial/purified non-cell systems or in cellular assays that poorly distinguish between multiple binding targets^31^ i.e. receptors and GAGs. It is therefore envisioned that these *in vitro* approaches will be broadly applicable to investigate secretion and binding of peptides and contribute to our understanding of chemokine-GAG binding and chemokine-mediated signalling in cells.

### Limitations of study

All experiments in this study were performed in model HEK293 cells therefore the broad applicability of the approach to other secreted peptides will depend on the ability to use CRISPR/Cas9 to edit cells of interest. In general, fusion of HiBiT to a protein may alter protein stability or expression and therefore the amount of HiBiT-tagged CXCL12 quantified may differ from the true levels of endogenous CXCL12 in HEK293 cells. Finally, it was not possible by luciferase complementation to determine to which specific types of proteoglycans/GAGs that CXCL12-HiBiT binds, but rather it was inferred that such interactions are occurring. Such specific interactions may be explored in the future using the NanoBRET binding approach described here.

## Resource availability

### Lead contact

Further information and requests for resources and reagents should be directed to and will be fulfilled by the Lead Contact, Stephen J Hill.

### Materials availability

Materials developed from this study are available from the Lead author on reasonable request.

### Data and code availability

This study did not generate datasets or code.

## Methods

All methods can be found in the accompanying Transparent Methods supplemental file.

## Acknowledgements

This work was supported by MRC grant number MR/N020081/1. C.W.W. is supported by an NHMRC CJ Martin Fellowship (1088334) and by a UWA fellowship support grant. We would like to thank Dr. Brigit Caspar for generating the constructs encoding SNAP/CXCR4.

## Author contributions

C.W.W and S.J.H conceived the study. C.W.W. designed the experiments, generated reagents, conducted the experiments and performed the data analysis. C.W.W., K.D.G.P. and S.J.H. wrote or contributed to the writing of the manuscript.

## Supplementary Information

### Transparent Methods

#### Materials

AMD3100 was purchased from Selleckchem (USA). Heparan sulfate (Hepran sulfate) sodium salt from bovine kidney and Surfen hydrate (Surfen) were from Sigma-Aldrich (United Kingdom). Furimazine, purified HiBiT (HiBiT-Halotag, control peptide) and purified LgBiT NLuc fragments were purchased from Promega (USA). Membrane impermeant SNAP-tag AF488 was purchased from New England Biolabs (United Kingdom). AMD3100 (10 mM) and heparan sulfate were dissolved in water. Surfen hydrate (10 mM) was dissolved in Dimethyl sulfoxide (DMSO). All further dilutions were performed in assay buffer containing 0.1% bovine serum albumin (BSA, Sigma-Aldrich, United Kingdom).

#### Molecular Biology

The CXCR4 cDNA sequences were provided through the ONCORNET consortium from Vrije Universiteit Amsterdam in pcDEF3 plasmids. pCDNA3.1 (+) neo expression constructs encoding SNAP/CXCR4 were generated as described previously (White et al., 2020), except that sig-SNAP (Gherbi et al., 2015) was ligated in frame using the restriction enzymes BamHI and XhoI in place of sig-NLuc.

#### CRISPR/Cas9 genome engineering

Guide RNA construction was performed as described previously in the detailed protocol (Ran et al., 2013). Briefly, guide sequences were designed using the CRISPR Design Tool (http://crispr.mit.edu/) to target the C-terminus of CXCL12 (ACTTGTTTAAAGCTTTCTCC) and ligated as complementary oligonucleotides 5’-AAACGGAGAAAGCTTTAAACAAGTC-3’ and 5’-CACCGACTTGTTTAAAGCTTTCTCC-3’ into the pSpCas9(BB)-2A-Puro (PX459 V2) expression construct (from Feng Zhang, Addgene plasmid # 62988) linearized by the restriction enzyme BbsI (NEB). To introduce DNA encoding GSSG-HiBiT into the CXCL12 genomic locus, a donor repair template was designed using the human genome assembly (GRCh38/hg38) and UCSC genome browser (http://genome.ucsc.edu/). The repair template was synthesised as single stranded oligo DNA nucleotides (ssODN, Integrated DNA Technologies, Inc. (IDT)) and consisted of homology arms surrounding GSSG-HiBiT with the CXCL12 stop codon deleted and the PAM motif mutated by a silent C-T substitution. The sequence used was 5’-GAACAACAACAGACAAGTGTGCATTGACCCGAAGCTAAAGTGGATTCAGGAGTATCTGG AGAAAGCTTTAAACAAGGGGAGTTCTGGCGTGAGCGGCTGGCGGCTGTTCAAGAAGATTAGCTAAGCACAACAGCCAAAAAGGACTTTCCGCTAGACCCACTCGAGGAAAACTAAAACCTTGTGAGAGATGAAAGGGCAAA-3’. Positive clones were genotyped using Q5® High-Fidelity DNA Polymerase (New England Biolabs, UK) as per the manufacturer’s instructions and the oligonucleotides 5’-CCTTCCTCCTGTGCAGCC-3’ and 5’-CAGGGTCTAAATGCTGGCAA-3’, which anneal outside the ssODN repair template.

#### Cell culture

HEK293T cells were maintained in Dulbecco’s Modified Eagle’s Medium (Sigma Aldrich) supplemented with 10% fetal calf serum at 37°C/5% CO_2_. Transfections were performed using FuGENE (Promega, USA) according to the manufacturer’s instructions. Cells were passaged or harvested using PBS (Sigma Aldrich) and trypsin (0.25% w/v in versene; Sigma Aldrich). CRISPR/Cas9 genome-engineering of HEK293 cells was performed as described previously (Ran et al., 2013; White et al., 2017). Briefly, HEK293T cells were seeded in 6 well plates at 300,000-400,000 cells per well and incubated for 24h at 37°C/5% CO_2_. Cells were then transfected with px459 sgRNA/Cas9 expression constructs and the ssODN donor repair template. Cells were cultured for 24h then treated with puromycin (0.3 µg/ml, Sigma-Aldrich) for 3 days to select for transfected cells. Following selection, cells were single cell cloned and allowed to expand for 2-3 weeks. Following expansion single clones expressing CXCL12-HiBiT were screened for luminescence following the addition of furimazine (10 µM) and purified LgBiT (10 nM) using a PHERAStar FS plate reader.

#### CXCL12-HiBiT assays

To investigate CXCL12-HiBiT secretion, wildtype or HEK293 cells expressing genome-edited CXCL12-HiBiT were seeded into poly-D-lysine coated white flat bottom 96 well plates at 30,000 or 60,000 cells/well and incubated for 24h at 37°C/5%CO_2_. On the day of the assay, cells were washed and incubated with pre-warmed 1x HEPES Buffered Salt Solution (1xHBSS; 25mM HEPES, 10mM glucose, 146mM NaCl, 5mM KCl, 1mM MgSO4, 2mM sodium pyruvate, 1.3mM CaCl2, 1.8g/L glucose; pH 7.2) supplemented with 0.1% BSA for 1, 2 or 4 hours or immediately incubated with purified LgBiT (30 nM) and furimazine (10 µM). Total luminescence was then measured on a PHERAStar FS plate reader. Analysis of the effect of time and cell number on CXCL12-HiBiT expression was performed at 20 minutes post LgBiT addition. In assays to investigate the effect of glycosaminoglycan or CXCR4 modulation on the levels of observable CXCL12-HiBiT, cells were washed with HBSS then incubated with HBSS supplemented with 0.1% BSA and 30 nM purified LgBiT for 2h 37°C. Furimazine (10 µM) was added to cells and allowed to equilibrate for 5 minutes before total luminescence was measured on a PHERAStar FS plate reader and 5 basal reads were taken. At time = 0, HBSS, AMD3100 (1 µM), surfen (10 µM), heparin sulfate (30 µg/mL), AMD3100 (1 µM) plus heparin sulfate (30 µg/mL) or surfen (10 µM) plus heparin sulfate (30 ug/mL) were added to the wells and total luminescence was measured. Baseline corrected luminescence was calculated by subtracting vehicle (HBSS)-treated luminescence from the ligand-treated luminescence.

#### Quantification of tagged protein by luciferase activity

To quantify CXCL12-HiBiT expression, wildtype HEK293 cells or HEK293 cells expressing genome-edited CXCL12-HiBiT were seeded into poly-D-lysine coated white flat bottom 96 well plates at 30,000 or 60,000 cells/well and incubated for 24h at 37°C/5% CO_2_. On the day of the assay, genome-edited CXCL12-HiBiT HEK293 cells were washed and incubated with pre-warmed HBSS for 2h at 37°C. A log HiBiT control protein (HiBiT-HaloTag, Promega, USA) standard curve (10 fmol - 100 nM) was constructed in parallel by diluting the purified HiBiT control protein in HBSS supplemented with 0.1% BSA and adding to wells containing wildtype HEK293 cells. Purified LgBiT (100 nM) was then added to each well and cells incubated for a further 5 minutes before 10 μM furimazine was added and total light emissions were measured on a PHERAStar FS plate reader.

#### CXCL12-HiBiT NanoBRET ligand binding

For CXCL12-HiBiT NanoBRET competition ligand binding assays, wildtype HEK293 cells were seeded in 6 well plates at 300,000 cells per well and incubated for 24h at 37°C/5% CO_2_. Cells were then transfected with 500 ng/well pcDNA3.1 (neo) plasmid encoding SNAP/CXCR4 and incubated for a further 24h. Transfected or un-transfected HEK293 cells were then seeded with HEK293 cells expressing genome-edited CXCL12-HiBiT into poly-D-lysine coated white flat bottom 96 well plates, at 20,000 cells/well of each cell line and incubated for 24h at 37°C/5% CO_2_. On the day of the assay, cells were incubated with 0.25 µM membrane impermeant SNAP-tag AF488 for 1h at 37°C/5% CO_2_ prepared in serum-free DMEM. After incubation, cells were washed 3 times with pre-warmed HBSS and incubated with purified 30 nM purified LgBiT in the absence or presence of AMD3100 (10 pM – 10 µM) for 2h at 37°C. Following ligand incubation, 10 μM furimazine was added and plates equilibrated for 5 mins at room temperature. Sequential filtered light emissions were recorded using a PHERAStar FS plate reader using 475 nm (30 nm bandpass) and 535 nm (30 nm bandpass) filters. BRET ratios were calculated by dividing the 535 nm emission (acceptor) by the 475 nm emission (donor).

#### Determination of CXCL12-HiBiT-LgBiT affinity

To investigate the affinity of CXCL12-HiBiT-LgBiT complementation, HEK293 cells expressing genome-edited CXCL12-HiBiT or wildtype HEK293 cells were seeded into poly-D-lysine coated white flat bottom 96 well plates at 30,000 cells/well and incubated for 24h at 37°C/5% CO_2_. On the day of the assay, cells were washed and incubated with HBSS supplemented with 0.1% BSA for 2hrs at 37°C. Cells expressing genome-edited CXCL12-HiBiT were then incubated with increasing concentrations of purified LgBiT for 30 minutes at 37°C in the absence or presence of Surfen (10 µM). In parallel, non-specific luminescence was determined by adding purified LgBiT to wells containing wildtype cells only. Following incubation, furimazine (10 µM) was added, plates incubated for 5 minutes, and total light emissions measured using a PHERAStar FS plate reader.

#### Data presentation and statistical analysis

BRET ratios were calculated by dividing the acceptor emission by the donor emission. Calculation of baseline corrected BRET ratios or luminescence values are described in the methods for each assay configuration.

Prism 7 software was used to analyse ligand-binding curves. For CXCL12-HiBiT – purified LgBiT saturation complementation assays, total and non-specific saturation binding curves were simultaneously fitted using the following equation:

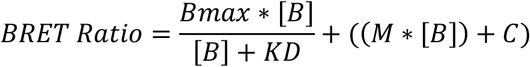

where Bmax is the maximal response, [B] is the concentration of LgBiT in nM, KD is the equilibrium dissociation constant in nM, M is the slope of the non-specific binding component and C is the intercept with the Y-axis. Luminescence generated by LgBiT [B] incubated on wildtype cells was used as non-specific binding.

Inhibition concentration response-data were fitted using the following equation:

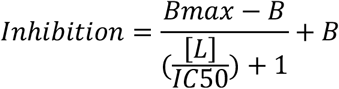

where Bmax is the maximum response of the probe, B is the non-specific binding or response, with both Bmax and B defined from the plateaus of the curve. [L] is the concentration of the competing ligand, IC50 is the concentration of the competition ligand required to inhibit 50% of the maximum response. pIC_50_ values were calculated as -log IC_50_.

Quantification of HiBiT-CXCL12 expression was interpolated by Prism from linear regression of a log-log standard curve fitted with the following equation:

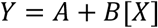

where [X] is the concentration of HiBiT, Y is the luminescence output, A is the y-intercept and B is the slope of the line. Complementation of HiBiT-LgBiT has been reported to produce a linear correlation extending over 8 orders of magnitude(Schwinn et al., 2018). Here we observed a linear correlation of R^2^ = 0.99, with a slope of 0.85 ± 0.04 (n=6). Statistical analysis was performed using Prism 7 software (GraphPad, San Diego, USA) using one or two-way ANOVA with appropriate multiple comparisons tests where required. Specific statistical tests used are indicated in the figure legends and were performed on the mean data of individual experiments (n) also indicated in the figure legends. A p-value <0.05 was considered statistically significant.

## Supplementary Information

**Supplementary Figure 1:**
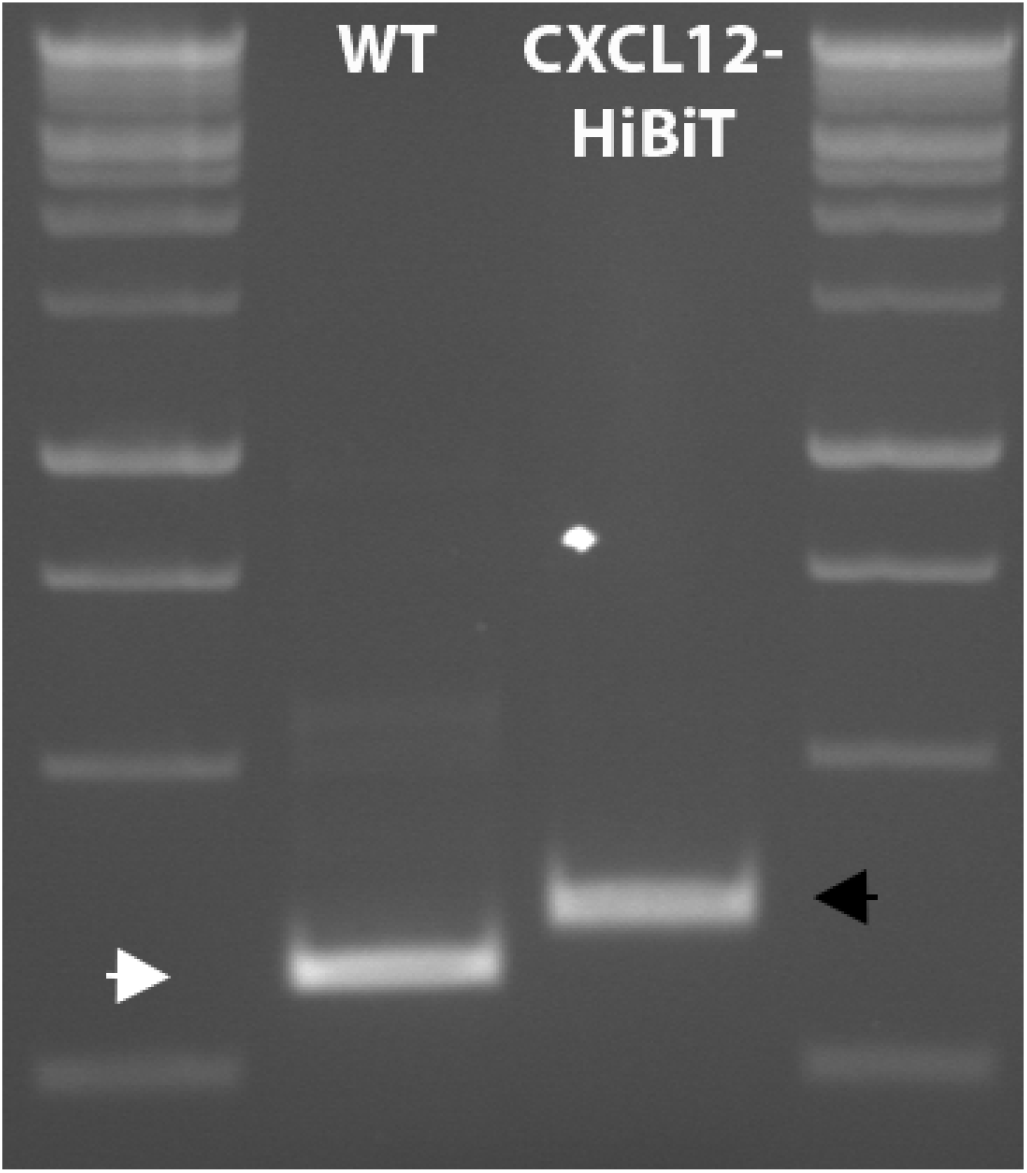
Genotyping of wildtype and genome-edited HEK293 cell lines. PCR amplification using primers targeting the CXCL12 locus of genomic DNA extracted from wildtype (WT) HEK293 cells (**lane 2)** or HEK293 cells genome-edited to express CXCL12-HiBiT (**lane 3)**. Wildtype PCR product at 314bp indicated by white arrow and black arrow indicates PCR product of inserted tag at 359bp. Lanes 1 and 4: Promega 1Kb benchtop DNA ladder.

